# Deterministic abiotic filtering and halophilic core microbiomes shape bacterial communities in costal salt flats (Sabkha) of southern Morocco

**DOI:** 10.64898/2026.01.12.698994

**Authors:** Ghita Amechatte, Nabil Radouane, Ayoub El Mouttaqi, Danilo Licastro, Abdelaziz Hirich, Hijri Mohamed, Bulbul Ahmed

## Abstract

Coastal salt flats, locally known as Sabkhas, are hypersaline, alkaline desert ecosystems that impose extreme abiotic stress on microbial and plant life. Despite their ecological significance, plant-associated microbiomes in these habitats remain poorly characterized. In this study, we investigated the bacterial communities of native halophytes across three sabkha sites in southern Morocco using an integrated culture-independent and culture-dependent framework. Soil physicochemical analyses revealed strong gradients in salinity and ionic composition, along with consistently alkaline pH across sites. These conditions strongly structured bacterial assemblage: alpha diversity declined progressively from bulk soil to rhizosphere soil, root and shoot; and beta diversity showed clear compartmental separation driven by environmental factors. Canonical Correspondence Analysis identified electrical conductivity (EC), Na₂O, K₂O and carbonate fractions as the main abiotic drivers. Across plant species, bacterial communities converged toward a stable halophilic core microbiome dominated by *Halomonas*, *Kushneria* and *Marinococcus*, with 66% of ASVs shared across compartments. Host identity played a secondary role, as environmental filtering overshadawed host-specific associations. Culture-dependent isolation recovered 19 halophilic and halotolerant bacterial strains, mainly, *Halomonas*, *Idiomarina*, *Marinobacter*, *Psychrobacter*, *Planomicrobium* and *Bac illus,* tolerating up to 25% NaCl. The strong concordance between cultured isolates and metabarcoding profile confirms that dominant halophilic lineages are both ecologically robust and readily culturable. Together, these findings demonstrate that sabkha plant microbiomes are primarily shalped by deterministic abiotic filtering and harbor resilient, stress-adapted bacterial communities. Sabkhas thus represent promising reservoirs of halophilic microbes with potential applications in saline agriculture and improving crop resilience under extreme environmental conditions.

**Importance:** Coastal salt flats (sabkhas) are among the most extreme terrestrial environments, characterized by high salinity, alkalinity, and limited water availability. As soil salinization expands worldwide, understanding how life persists in such habitats is increasingly important for sustainable agriculture. This study shows that sabkha ecosystems impose strong environmental filtering on plant-associated bacterial communities, leading to highly structured microbiomes across soil, root, and shoot compartments. Despite differences among sites and plant species, bacterial communities converged toward a shared halophilic core microbiome, dominated by salt-adapted genera that are resilient to extreme ionic stress. Importantly, many of these dominant bacteria were readily culturable, highlighting sabkhas as accessible reservoirs of stress-tolerant microbes. Our findings demonstrate that abiotic conditions outweigh plant identity in shaping microbiome assembly under extreme stress and reveal sabkha halophytes as valuable natural models for discovering microbes with potential applications in saline agriculture, soil restoration, and crop resilience in salt-affected environments.

## Introduction

Sabkha ecosystems provide a compelling natural context for studying microbial adaptation to extreme abiotic stress. The term Sabkha, derived from the Arabic word for “salt flat,” refers to hypersaline landscapes distributed across North Africa, the Arabian Peninsula, and parts of Australia. Both coastal and inland sabkhas experience extreme salinity, high temperatures, low water availability, and intense evaporation, resulting in soils with elevated pH, high carbonate content, and distinctive ionic profiles [1]. Similar environmental constraints have been shown to shape the root-associated microbiomes of plants in Sahara desert in the southern Morocco, where both soil physicochemical properties and plant identity influence bacterial community composition [2]. Despite these harsh abiotic conditions, sabkhas host specialized halophytic vegetation whose associated microbiomes remain largely unexplored.

Increasing evidence highlights the central role of environmental context in shaping plant-microbe interactions. Environmental factors, such as soil nutrients, pH, temperature, salinity, and water availability, are major determinants of the composition and function of plant-associated microbial communities [3–6]. In benign and moderate environments, plants often exert strong control over microbiome assembly through root exudates, immune signaling and developmental cues, enabling them to recruit beneficial microbes and structure highly diverse, host-specific communities [6, 7]. In contrast, in extreme ecosystems, such as saline soils, arid regions, or sabkhas, abiotic constraints including high salinity, drought, or elevated temperatures frequently override host-driven selection, limiting microbial diversity and narrowing microbial plasticity. These stressful conditions often lead to the convergence of taxonomically unrelated plants toward functionally similar or stress-tolerant microbial communities enriched in halophilic or extremotolerant taxa [5, 8–10]. Transitional or disturbed environments may show intermediate patterns in which both host and environmental drivers operate in context-dependent ways, producing dynamic shifts in microbial richness, functional potential, and community resilience [11–13].

These contrasting mechanisms have important implications for climate change and agricultural adaptation. In harsh or rapidly changing environments, strong environmental filtering can reduce the plant’s ability to shape its microbiome; however, the persistence of functionally redundant, stress-tolerant microbial groups may still confer essential support for plant survival [4, 8, 14]. Leveraging beneficial microbes, including halotolerant bacteria and stress-adapted mycorrhizal fungi, represents a promising strategy to enhance crop performance in degraded or saline soils, provided that microbial traits are well matched to local abiotic conditions [2, 10, 14, 15]. Soil salinity imposes severe stress on plants but can be mitigated through inoculation with salt-tolerant rhizobacteria that improve plant salt resistance and growth in saline soils. Halotolerant plant growth promoting rhizobacteria have been shown to help plants tolerate salinity stress and support sustainable agriculture in salt-affected soils. Therefore, understanding how host genetics interacts with environmental context is fundamental for predicting microbiome assembly under climate change and for designing effective microbial-based interventions [6, 16, 17]. Collectively, these insights reinforce that environmental context is a primary determinant of plant–microbe interactions, particularly under extreme abiotic stress where environmental filtering can dominate over host control.

Plants coexist with diverse microbial communities that colonize both external and internal tissues, forming complex holobionts essential for plant function and fitness. Among these microbial partners, bacteria are particularly influential due to their abundance, metabolic versatility, and capacity to form intimate associations with a wide range of plant hosts. The assembly of plant-associated microbiomes is shaped by a combination of host-derived factors, including genotype, root exudation patterns, developmental stage, and immune activity, and environmental factors [18, 19]. Life-history traits also influence microbial recruitment; for example, perennial plants often harbor richer and more phylogenetically diverse microbiomes than annuals, owing to their longer lifespans, deeper root systems, and more stable rhizosphere environments [20]. Yet, under extreme abiotic stress, such as high salinity, drought, alkalinity, or heat, soil physicochemical properties frequently emerge as the dominant determinants of microbial community structure [21, 22]. Across these ecosystems, taxonomically unrelated plants may converge on similar microbiomes enriched in stress-tolerant bacterial lineages such as, *Halomonas*, *Marinobacter*, *Salinimicrobium* and *Pseudomonas,* reflecting environmental determinism rather than host specificity [23]. These patterns underscore the need to disentangle the relative contributions of host-driven and environment-driven selection in microbiome assembly.

In this study, we investigated the structure and assembly of microbiomes associated with native halophytes at three sites within the inland Oum Dbaa sabkhas in southern Morocco. Using integrated culture-independent and culture-dependent approaches, we aimed to determine whether microbiome assembly in these extreme habitats is governed predominantly by abiotic filtering or by host-specific selection. We hypothesized that: (i) strong edaphic constraints, particularly salinity, ionic composition, and alkalinity, act as the primary drivers of bacterial community structure, overriding host genotype effects; (ii) bacterial communities converge toward a shared halophilic core microbiome across different plant species, indicative of deterministic environmental filtering; and (iii) the dominant halophilic taxa detected through metabarcoding will also be recoverable through cultivation, reflecting their ecological robustness and applied potential. By combining community profiling with culturable strain isolation, this study provides a comprehensive mapping of bacterial communities in sabkha ecosystems and identifies promising halophilic and halotolerant candidates for developing stress-resilient microbial inoculants to support sustainable agriculture under extreme environmental conditions.

## Methods

### Study site and sampling

Sampling was conducted at three sites within the Oum Dbaa inland sabkha ecosystem in southern Morocco (Fig.1 A). The study sites were located at 27°27′27″N, 13°02′59″W; 27°27′12″N, 13°02′42″W; and 27°26′46″N, 13°01′58″W. The geographical map of sampling locations was generated in R (v4.3.1) using the packages ggplot [24], sf [25] and rnaturalearth [26], based on the recorded GPS coordinates of each site. At each sabkha, representative halophytic plant species were sampled for microbiome analysis (Table S1). A total of ten halophytes were included (Fig. 1C-D), encompassing shrubs (*Arthrocaulon macrostachyum* (Moric.) Piirainen & G. Kadereit, *Halocnemum strobilaceum* (Pall.) M. Bieb), small trees (*Tamarix amplexicaulis* Ehrenb., *Nitraria retusa* (Forssk.) Asch.), grasses (*Juncus acutus* L.), and perennial herbs (*Saharanthus ifniensis* (Caball.) M. B. Crespo & Lledó, *Caroxylon tetrandrum* (Forssk.) Akhani & Roalson, *Caroxylon tetragonum* (Dalile) Moq., *Tetraena gaetula* (Emb. & Maire) Beier & Thulin., *Frankenia corymbosa* Desf.). For each plant species, four biological replicates were collected, and from each replicate, three compartments were sampled: rhizosphere soil, root, and shoot tissues (Fig. 1B). At each site, three bulk soil samples (0-20 cm depth) were collected away from plant roots to serve as environmental controls and for soil physicochemical properties. In total 151 samples were obtained, including 142 plant-associated samples and 9 bulk soil samples. Immediately after collection, all samples were stored in a refrigerator at ∼4° C in the mobile field laboratory vehicle (African Sustainable Agriculture Research Institute - ASARI, Laayoune, Morocco), then transported and stored at –20° C prior to DNA extraction and isolation of bacteria.

**Figure 1.**
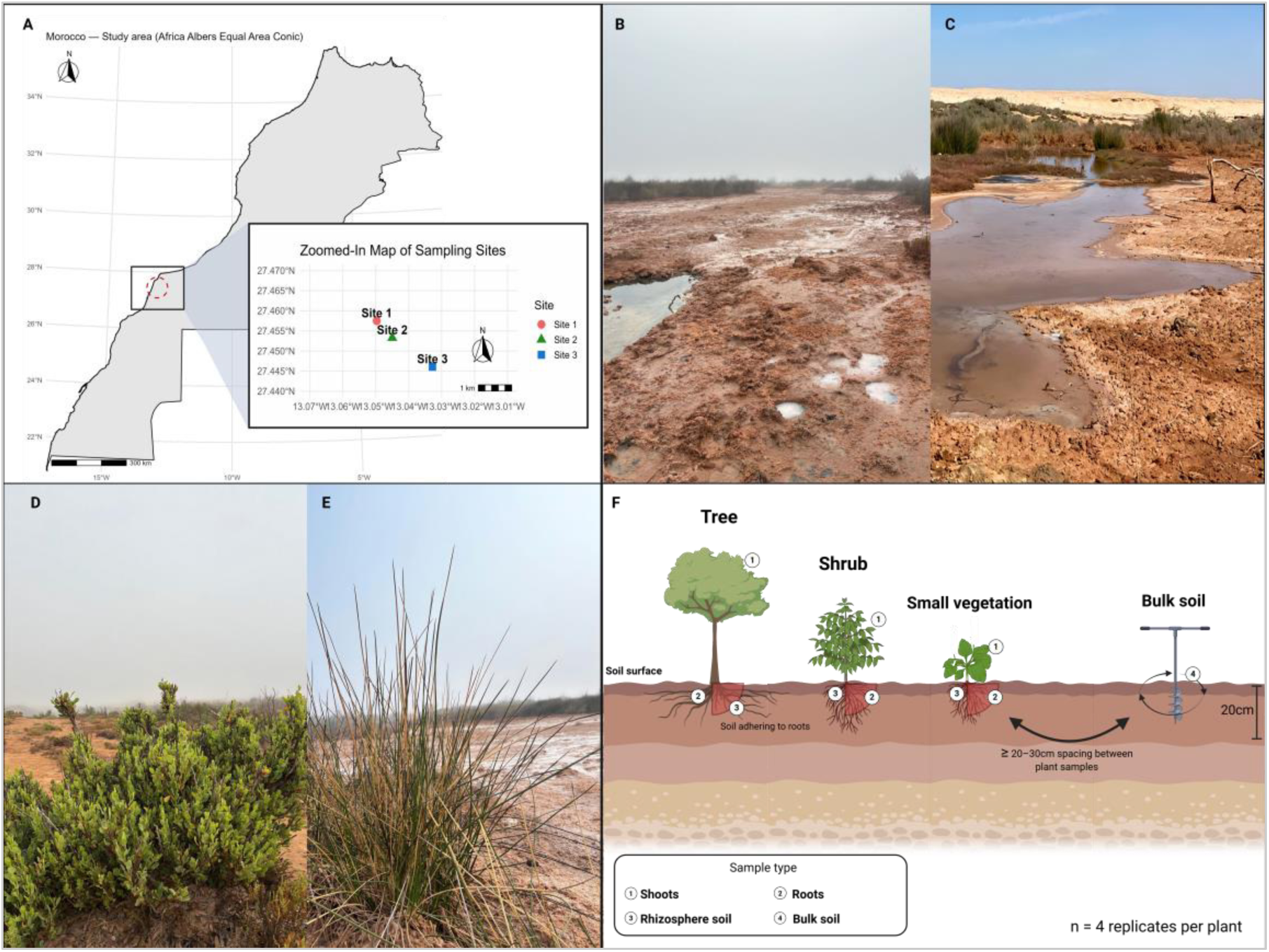
Study areas, sampling strategy and sabkha landscape. (A) Map showing the three sampling sites within sabkha ecosystems of Southern Morocco. Inset shows a zoomed view of the spatial distribution of the three sites. (B-C) Field of the sabkha environments. (D-E) dominant vegetation types sampled in this study, including examples of typical saline terrain and halophytic plant species. (F) Schematic illustration of the sampling design across plant functional groups: trees, shrubs and small vegetation. For each plant, four compartments were sampled: shoot ①, root ②, rhizosphere soil ③, and bulk soil ④, with ≥ 20-30 cm spacing between replicates (n = 4 per plant type).

### Soil physicochemical analysis

Soil physicochemical analyses were performed characterization of soils was carried out at the ASARI (Laayoune, Morocco). Each soil replicate was analyzed individually across all parameters, without homogenization, to reflect natural site-level variability. Soil pH was measured in a 1:2.5 soil-to-water suspension following ISO 10390:2005. Electrical conductivity (EC) was determined in the saturated paste extracts according to NF ISO 11265, and soil salinity was inferred from EC values. Total carbonate content was quantified by the ISO 10693 (1995) volumetric method, and active carbonate by NF X 31-106. Available phosphorus (P₂O₅) was measured using the Olsen extraction method (NF ISO 11263). Organic carbon (OC) was analyzed following NF ISO 14235 and converted to organic matter (OM) using OM = OC × 1.724. Total nitrogen (N) was measured using the Kjeldahl digestion method. For elemental composition, soil samples were digested with concentrated sulfuric acid, and macroelements (K₂O, CaO, Na₂O, MgO) and microelements (Fe, Zn, Cu, Mn) were quantified by atomic absorption spectroscopy (AAS).

### Bacterial isolation

Bacteria isolation from the root, shoot, and rhizosphere samples followed the protocol of [27]. Plant tissues were rinsed with sterile saline solution to remove surface debrits, cut into ∼1 cm fragments, and placed onto Marine Agar (Difco™ 2216; BD Difco, Franklin Lakes, NJ, USA) plates to enable the growth of endophytic bacteria. For rhizosphere samples, approximately 1 g of soil was suspended in 9 mL of sterile saline solution and shaken at 150 rpm for 45 min to detach microbial cells [28]. Suspension was serially diluted (10^-^¹-10^-^⁶), and pilot tests determined the optimal dilution for plating. Aliquots (100 μL) were spread onto Marine Agar plates and incubated at 28°C. Colony emergence generally occurred after 48-72 hours, though some isolates required 96 to 120 hours. Distinct colonies were repeatedly re-streaked to obtain pure isolates. A single colony from each isolate was subjected to direct colony PCR for taxonomic identification. One colony was transferred into 10 µL sterile nuclease-free water, macerated and heated at 98 °C for 10 min. The resulting lysate served as PCR template. Amplification of the 16S rRNA gene used primers 27F (AGAGTTTGATCCTGGCTCAG) and 1492R (TACGGHTACCTTGTTCGACTT). PCR reactions (50 µL) contained 2 µL of each primer (10 µM), 10 µL GC enhancer reagent, 25 µL Master Mix (New England Biolabs, Ipswich, MA, USA), 1 µL of colony lysate, and nuclease-free water. Cycling conditions were: 95 °C for 5 min; 35 cycles of 95 °C for 30 s, 57 °C for 60 s, and 72 °C for 60 s; followed by final extension at 72 °C for 10 min. Amplicons were visualized on 1% agarose gels using an iBright imaging system (Thermo Fisher Scientific, Waltham, MA, USA). Successful PCR products were purified and subjected to Sanger sequencing at Genome Quebec (Montréal, Canada). Forward and reverse sequences were assembled, quality-trimmed, and merged into contig using Geneious Prime® v2025.2.1 (Biomatters, Auckland, New Zealand). Taxonomic identification was performed using BLASTn against the NCBI nucleotide database; only hits with ≥ 97% sequence identity and ≥ 98% coverage were accepted for genus- or species-level classification.

### DNA extraction, library preparation and sequencing

Total genomic DNA was extracted using previously established methods [29] from 250 mg of rhizosphere and bulk soil using the DNeasy PowerSoil Pro Kit (Qiagen, Toronto, ON, Canada) and from 100 mg of grounded root tissue using the DNeasy Plant Mini Kit (Qiagen, Toronto, Canada), following the manufacturer’s instructions. Prior to homogenization, plant samples were flash-frozen in liquid nitrogen, while soil samples were processed directly. Both soil and plant samples were homogenized using a TissueLyser II (Qiagen, Hilden, Germany) with 2 mm tungsten beads at 24 Hz for 15 min. DNA was eluted in 50 µL of elution buffer for soil samples and in 20 µL for plant samples and stored at -20 °C until further use. Extracted DNA concentration and purity were assessed using a BioSpectrophotometer (Eppendorf, Hamburg, Germany) and a NanoDrop ONE spectrophotometer (Thermo Scientific, Wilmington, DE, USA) and DNA integrity were verified by electrophoresis on 1% agarose gels stained with GelRed (1/10,000) and visualized using the GelDoc system (Bio-Rad, Montreal, QC, Canada). For bacterial community profiling, the V3-V4 region of the 16S rRNA gene was amplified with primers CS1_341 (5′-ACACTGACGACATGGTTCTACACCTACGGGNGGCWGCAG-3′) and CS2_806R (5′-TACGGTAGCAGAGACTTGGTCTGACTACHVGGGTATCTAATCC-3′).Libraries were prepared using the TruSeq Nano DNA LT Sample Preparation Kit (Illumina, United States) and sequenced on the Illumina NovaSeq 6000 platform with 2 × 150 bp paired-end reads at the Next Generation Sequencing facilities of LAGE (Laboratory of Genomics and Epigenomics) in Area Science Park, Trieste, Italy.

### Bioinformatic analysis of 16S rRNA sequences

Raw paired-end NovaSeq reads were processed in R v4.0.0 using the DADA2 package v1.18.0 [30]. Quality filtering discarded reads containing ambiguous bases and those with more than three expected errors (maxEE = 3). No read truncation was applied, and PhiX spike-in contamination was removed. Denoising was performed with the DADA algorithm, followed by merging of paired reads and removal of chimeric sequences using the consensus method. Because NovaSeq quality scores are binned, which can lead to spurious sequence variants [31], only amplicon sequence variants (ASVs) present at least 250 times across the dataset were retained for downstream analysis. Taxonomic assignment was carried out with the assign Taxonomy function in DADA2 against the SILVA database (release 138.2) [32] trimmed to the sequenced region. Non-bacterial sequences (chloroplast, mitochondrial) were removed prior to downstream analyses.

### Statistical analysis

To evaluate the effects of host identity and environmental variables on bacterial community structure, we conducted permutational multivariate analyses of variance (PERMNOVA) using the adonis2 function in the vegan R package (v2.6-4) [33]. Bray-Curtis dissimilarities were computed from genus-level relative abundance data after prevalence filtering. Each factor was first tested independently in univariable PERMANOVAs using 999 permutations and resulting p-values were adjusted for multiple testing with the Benjamini-Hochberg false discovery rate (FDR). To limit the influence of multicollinearity, variables with pairwise correlation coefficients |r| > 0.99 were excluded. The remaining predictors (n = 12) were included in a multivariable PERMANOVA, and the unique contribution of each variable was quantified by marginal R² values.

To correct for differences in sequencing depth, ASV counts were normalized to relative abundances using the “compositional” transformation implemented in the phyloseq package [34]. Bacterial alpha diversity was quantified using the Shannon, Simpson, and Chao1 indices via the estimate_richness function in phyloseq. Group differences were tested with the Kruskal-Wallis test, followed by pairwise Wilcoxon rank-sum tests adjusted with Bonferroni correction. Boxplots were generated using ggplot2 [24], with statistical annotations added through ggpubr [35].

Beta diversity was assessed using Bray-Curtis dissimilarities derived from ASV-level relative abundances. Group-level differences were tested by PERMANOVA (adonis, vegan), and the homogeneity of multivariate dispersions among groups was evaluated using the betadisper function in the same package. To identify microbial taxa differentiating plant compartments, we combined Random Forest classification and Indicator Species Analysis (IndVal.g). Random Forest models were trained using the randomForest R package [36] to rank genera by permutation importance, whereas IndVal.g (indicspecies package; [37] identified compartment-specific indicators based on permutation testing with FDR correction. These complementary approaches jointly revealed microbial genera that significantly contributed to the differentiation of bacterial communities of bulk soil, rhizosphere, root and shoot. Detrended correspondence analysis (DCA) [38]was first applied to the ASV abundance matrix to assess the length of underlying ecological gradients and to guide the selection of an appropriate constrained ordination method. Gradient lengths were evaluated in standard deviation units along the DCA axes. Because the length of the first DCA axis exceeded 4 standard deviation units, indicating unimodal species responses, canonical correspondence analysis (CCA) [39] was subsequently used to examine relationships between bacterial community composition and environmental variables.

## Results

### Soil physicochemical properties in the rhizosphere soils reflected bulk soil salinity gradients in sabkha

Comprehensive soil profiling across the three sabkha sites revealed pronounced heterogeneity in physicochemical conditions, with strong gradients in salinity and ionic composition (Fig. S1; Table S2). Site 1 exhibited the highest electrical conductivity (EC) in bulk soil, consistently exceeding 19,000 µS cm⁻¹, whereas Sites 2 and 3 showed substantially lower EC values (< 7,000 µS cm⁻¹) (Kruskal–Wallis, p < 0.05). Sodium oxide (Na₂O) and potassium oxide (K₂O) levels mirrored this pattern, reaching up to 51,916 mg kg⁻¹ Na₂O and 36,877 mg kg⁻¹ K₂O in Site 1. Despite differences in salinity, all soils maintained uniformly alkaline pH (8.4-9.2) and high carbonate content (18–36% CaCO₃), reflecting the inherently harsh baseline conditions of sabkha ecosystems. To evaluate how these edaphic differences translate to the plant microenvironment, we analyzed physicochemical properties of rhizosphere soil (Fig. S2). EC, Na₂O and K₂O remained significantly higher in Site 1 compared with Sites 2 and 3 (Kruskal-Wallis with Dunn’s test, p < 0.05), confirming that plants in Site 1 experience extreme ionic stress, whereas plants in Site 3 are exposed to comparatively milder conditions.

### Alpha- and beta-diversity analyses reveal strong soil**–**plant filtering

Sequencing of the 16S rRNA gene dataset yielded 6,010 bacterial ASVs across 151 samples. Taxonomic classification resolved 6,010 ASVs at the phylum level, 6001 at the order level and 2,316 at the genus level. Sequencing depth varied widely (9,820 - 671,482 reads per sample; mean = 228,014; median = 181,978). Rarefaction curves indicated sufficient sequencing coverage across all sample types (Fig. S3).

Alpha diversity showed clear influence of site, plant compartment, and to lesser extent plant type (Fig. 2 A-C; and Fig. S3). Chao1 richness was significantly lower at Site 1 than Sites 2 and 3 (p < 0.05), while Shannon and Simpson diversity did not differ significantly among sites (Fig. S4). Diversity did not vary significantly across plant types (trees, shrubs, small vegetation) (Fig. S4 D-F and Table S4), indicating limited influence of host growth form on richness or evenness. In contrast, sample types (Table S5) strongly structured microbial diversity. All three indices (Chao1, Shannon, and Simpson) declined along the soil-plant continuum, with bulk soil showing the highest richess and eveness, followed by rhizosphere, roots, and shoots. Significant reductions occurred between bulk soil and both rhizosphere (Chao1: *p* = 0.039; Shannon and Simpson: *p* = 0.005) and roots (Chao1: *p* = 0.010; Shannon and Simpson: *p* < 0.001). Diversity decreased further from roots to shoots (Chao1: *p* = 0.005; Shannon and Simpson: *p* = 0.001) (Fig. 2 A-C and Table S5). Differences between rhizosphere and plant tissues were weaker and non-significant, indicating that the strongest filtering steps occur during transitions into the plant and from roots to aboveground tissues.

**Figure 2.**
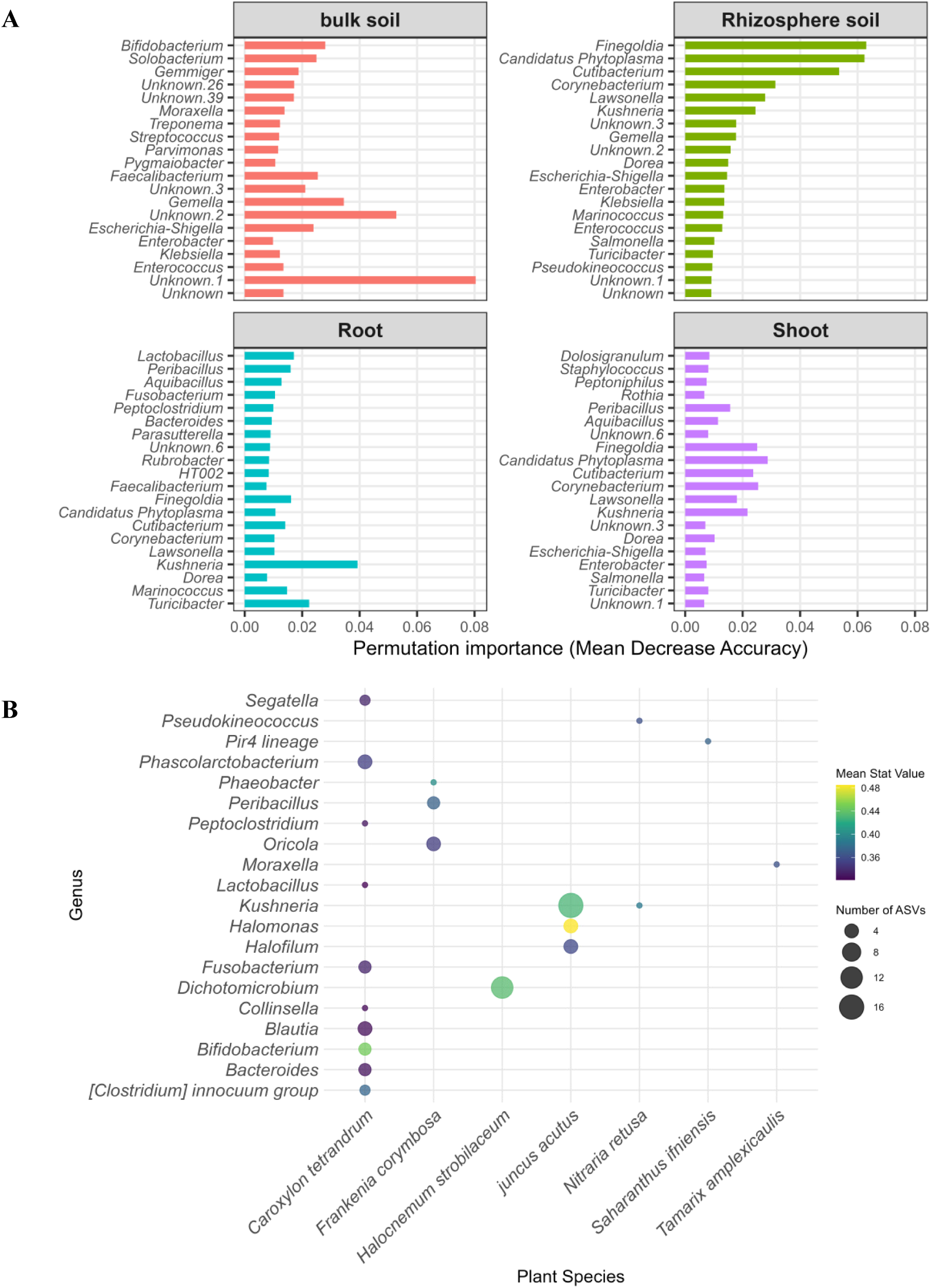
Alpha and beta diversity of bacterial communities across sample compartments. Boxplots shows alpha diversity by sample type based on (A) Chao1 richness, (B) Shannon diversity and (C) Simpson index. P-values indicated results of Kruskal-Wallis tests with Bonferroni-corrected pairwise comparisons. (D-F) Principal coordinates analysis (PCoA) of Bray-Curtis dissimilarities reveals differences in community composition. (D) sample type, (E) site, and (F) plant type. PERMANOVA p-values and homogeneity of dispersion results are shown for each panel.

Beta-diversity analyses (Bray-Curtis) were consistent with these patterns and supported by PERMANOVA results across all factors (Fig. 2D-F; Table S6). Microbial communities differed significantly by sample type (*p* = 0.001; Fig. 2D), by site (*p* = 0.09; Fig. 2E), and by plant type (*p* = 0.006; Fig. 2F). Beta-dispersion tests were also significant for each factor (*p* = 0.001), indicating that both differences in centroid positions and variability among groups contributed to the observed patterns (Table S6). Overall, these results suggest that environmental filtering along the soil–plant continuum is the dominant force shaping microbiome differentiation, while site-level heterogeneity and host identity exert secondary influences.

### Bacterial diversity and richness respond to spacial distribution

Taxonomic profiling revealed clear structuring of bacterial communities across compartments (Fig. 3). At the phylum level (Fig. 3A), microbiomes were dominated by Pseudomonadota, followed by Bacillota and Cyanobacteriota with pronouced compartment-specific differences. Bulk soil was enriched in Pseudomonadota, followed by Bacteroidota. The rhizosphere soil was enriched with Cyanobacteriota, followed by Bacillota and Pseudomonadota. Root and shoot tissues showed reduced phyla-level diversity but increased dominance of Pseudomonadota and Cyanobacteriota. At the genus level (Fig. 3B), bulk soil and rhizosphere compartments displayed broad taxonomic richness, including *Escherichia Shigella, Enterococcus, and Enterobacter*. In contrast, rhizosphere soil, root and shoot compartments were dominated by unclassified taxa. Root and shoot were also dominated by specialized halotolerant genera, such as *Kushneria* sp., *Marinococcus* sp., and *Halomonas* sp., reflecting selective colonization of internal plant tissues. Although detailed inter-site comparisons were limited, rhizosphere communities appeared more diverse and compositionally distinct than bulk soil, with potential site-specific enrichment of certain genera (e.g., *Blautia*, *Gemella*, *Peptoclostridium*). Further resolution of site-level variation may reveal subtle host or edaphic influences within each sabkha.

**Figure 3.**
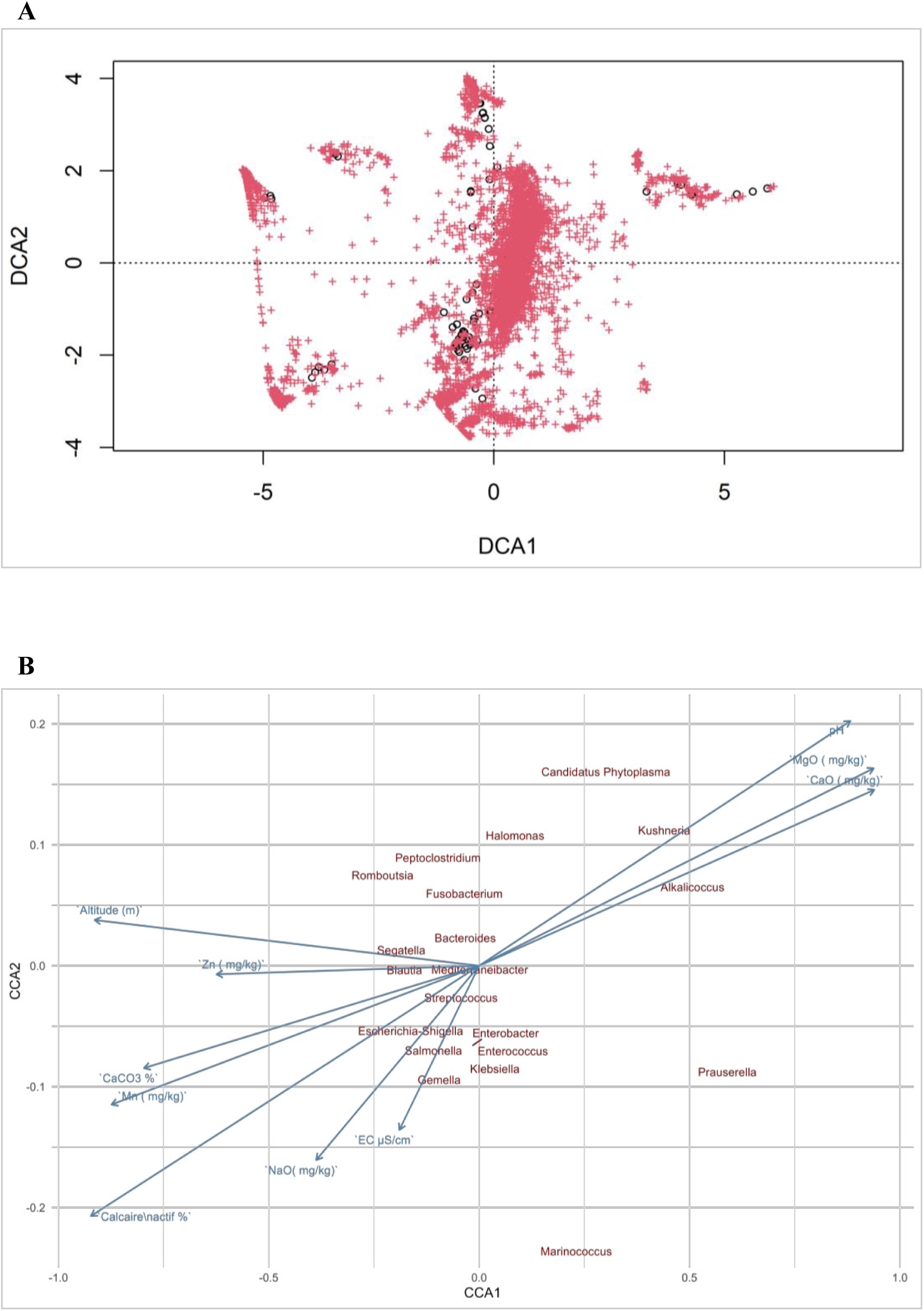
Relative abundance of dominant bacterial genera by sample type. (A) Mean relative abundance of dominant bacterial phyla across bulk soil, rhizosphere soil, root and shoot samples. (B) Mean relative abundance of dominant genera within each compartment. Only the most abundant taxa are shown; low-abundance genera are grouped as “Other”.

### Indicator bacterial species reveal strong compartment- and host-specific signatures

The analysis of bacterial indicator taxa across the three sabkhas revealed both distinct and overlapping microbial signatures among plant compartments and hosts (Fig. 4). Random Forest analysis identified clear compartment-specific patterns in bacterial community structure (Fig. 4A). Bulk soil communities were primarily characterized by several unclassified taxa, alongside genera such as *Bifidobacterium, Solobacteriu*m and *Gemella.* The rhizosphere was enriched in *Finegoldia*, *Candidatus Phytoplasma*, *Cutibacterium*, *Corynebacterium*, and several unclassified genera. Root tissues showed strong enrichment of *Kushneria*, *Lactobacillus*, and *Turibibacter*, reflecting strong selective filtering within the endosphere. Shoot-associated communities were characterized by *Candidatus Phytoplasma*, *Finegoldia*, *Cutibacterium*, *Corynebacterium*, *Kushneria*, and *Peribacillus*. These indicator genera, which achieved high permutation importance scores, demonstrate pronounced spatial partitioning along the soil–root–shoot continuum, highlighting the distinct ecological pressures structuring each plant compartment.

**Figure 4.**
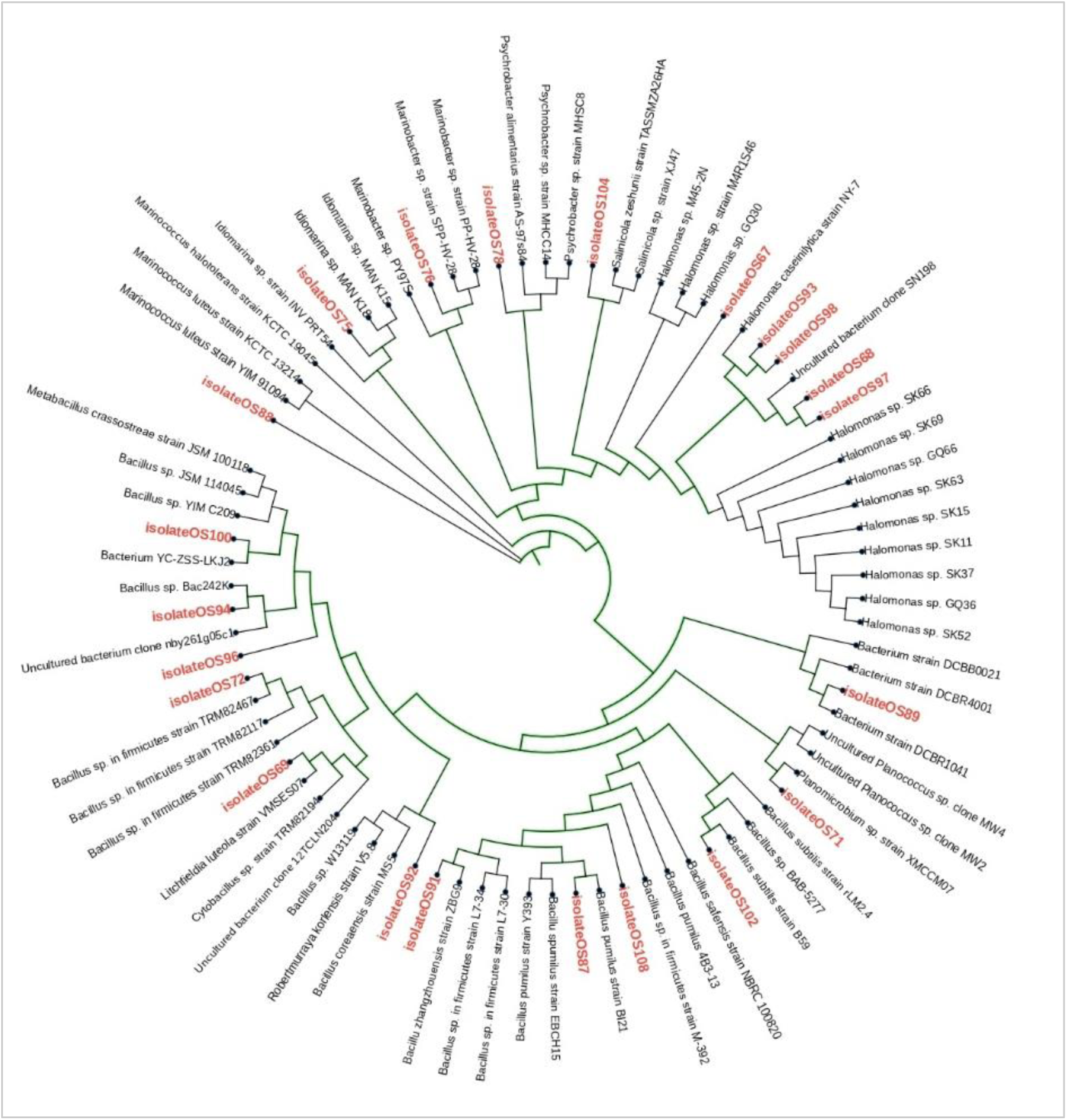
Random Forest and indicator species analyses. (A) Top 20 bacterial genera contributing to classification accuracy across four compartments (bulk soil, rhizosphere soil, root and shoot), ranked by permutation importance using Random Forest. (B) Indicator genera identified by IndVal analysis across plant species. Dot size corresponds to the number of ASVs, and color indicates the mean IndVal score (indicator strength).

To complement these patterns (Fig. 4A), we performed IndVal analysis to assess host-specific associations (Fig. 4B). *Juncus acutus* showed the strongest selective filtering, with high indicator values for the halophilic genera *Halomonas sp.* (IndVal = 0.492, *p* = 0.0004*), Kushneria sp.* (0.452, *p* = 0.0013), and *Halofilum sp.* (0.376, *p* = 0.0099). *Halocnemum strobilaceum* exhibited a single but highly robust indicator genus*, Dichotomicrobium* (IndVal = 0.448, *p* = 0.0018), consistent with its adaptation to extreme salinity and nutrient-poor soils*. Frankenia corymbosa* plant was enriched in *Phaeobacter sp.* (IndVal = 0.401, *p* = 0.0056), *Peribacillus* sp. (0.396, *p* = 0.0077), and *Oricola* sp. (0.357, *p* = 0.0245), and *Nitraria retusa* hosted two distinct indicators, *Kushneria* sp. (IndVal = 0.387, *p* = 0.0125) and *Pseudokineococcuss sp.* (0.362, *p* = 0.0148), whereas, *Saharanthus ifniensis* and *Tamarix amplexicaulis*, each displayed a single association, for instance, Pir4 lineage (IndVal = 0.374, *p* = 0.0107) and *Moraxella sp.* (IndVal = 0.363, *p* = 0.0134) respectively. *Caroxylon tetrandrum*, hosted the most diverse (10 genera) indicator profile, including *Bifidobacterium* sp. (IndVal = 0.469, *p* = 0.0002), *Clostridium innocuum* group (0.379, *p* = 0.0071), *Phascolarctobacterium sp.* (0.377, *p* = 0.0047), *Fusobacterium* sp. (0.356, *p* = 0.0113), *Segatella* sp. (0.355, *p* = 0.0126), *Bacteroides sp., Peptoclostridium sp., Blautia sp., Collinsella sp.,* and *Lactobacillus sp.* (Fig. 4B). Together, these results indicate that while environmental filtering dominates microbiome assembly, host identity modulates the specificity and breadth of microbial recruitment under extreme sabkha conditions.

### Core bacteriome and cross-host convergence

To identify conserved bacterial taxa across plant functional types (shrubs, small vegetation, and trees), we profiled the core genera within the shoot, root and rhizosphere compartments (Fig. S5 A-C) and assessed ASV-level diversity to evaluate cross-compartment microbial convergence (Fig. S5D). In the shoot compartment (Fig. S5A), *Kushneria* sp*., Enterobacter* sp*., Escherichia-Shigella* sp. and *Marinococcus* sp., were consistently detected across all plant types, with small vegetation exhibiting the highest core genus richness and relative abundance. Shrub shoots displayed slightly lower diversity, suggesting stronger selective pressures in aboveground tissues of woody perennials. Root-associated bacterial communities (Fig. S5B) showed a comparable core structure with dominated *Fusobacterium* sp.*, Enterococcus* sp, and *Peptoclostridium* sp. Tree roots harbored the richest and most evenly distributed core bacteriome, potentially reflecting more stable or complex rhizosphere–root interactions associated with perennial root systems. In rhizosphere soil (Fig. S5C), broader taxonomic diversity was observed, with *Escherichia*–*Shigella, Enterobacter* sp. and *Gemella* sp. dominating across all plant types. Trees again exhibited highed genus richness and compositional complexity than shrubs and small vegetation, consistent with increasing structured rhizosphere environments beneath long-lived plant canopies.

To evaluate community overlap, we quantified ASV-level intersections across compartments (Fig. S5D). A substantial fraction of ASVs (66%) was shared among roots, shoots, and rhizosphere soils, indicating the presence of a conserved microbial backbone spanning both rhizosphere and endophytic niches. Compartment-specific ASVs were most prevalent in roots (14%) and rare in shoots (1%), whereas the rhizosphere contained no unique ASVs, supporting its role as a microbiome reservoir seeding plant-associated communities. Together, these results highlight a conserved set of core bacterial genera across sabkha halophytes, with trees hosting the most diverse core microbiota and roots contributing most of the compartment-specific diversity.

### Environmental drivers of community structure

To elucidate the environmental factors shaping bacterial community composition in the sabkhas, we first assessed the underlying species response model using detrended correspondence analysis (DCA). The length of the first DCA axis was high (10.89 SD units), indicating strong species turnover and unimodal responses to environmental gradients (Fig. 5A; Table S7), and thereby justifying the use of Canonical Correspondence Analysis (CCA). The broad dispersion of taxa along DCA1 further suggests pronounced ecological differentiation among bacterial assemblages across sabkha environments, consistent with strong environmental filtering. Subsequently, CCA revealed clear relationships between dominant bacterial genera and soil physicochemical properties across sabkha environments (Fig. 5B; Table S8). Salinity, alkalinity, and calcareousness emerged as the strongest environmental correlates influencing bacterial community composition across sabkha soils (Table S8). Electrical conductivity (EC), soil pH, and both total and active carbonate fractions were significantly associated with community structure (*p* = 0.001). Genera such as *Kushneria, Halomonas*, and *Marinococcus* were strongly aligned oriented with high EC and carbonate gradients, reflecting their prevalence in saline-alkaline soils. In contrast, *Escherichia-Shigella and Enterococcus* were associated with nutrient-rich contexts under high EC conditions. Together, these results confirm that abiotic filters imposed by soil physicochemistry are the dominant drivers of sabkha bacteriome assembly.

**Figure 5.**
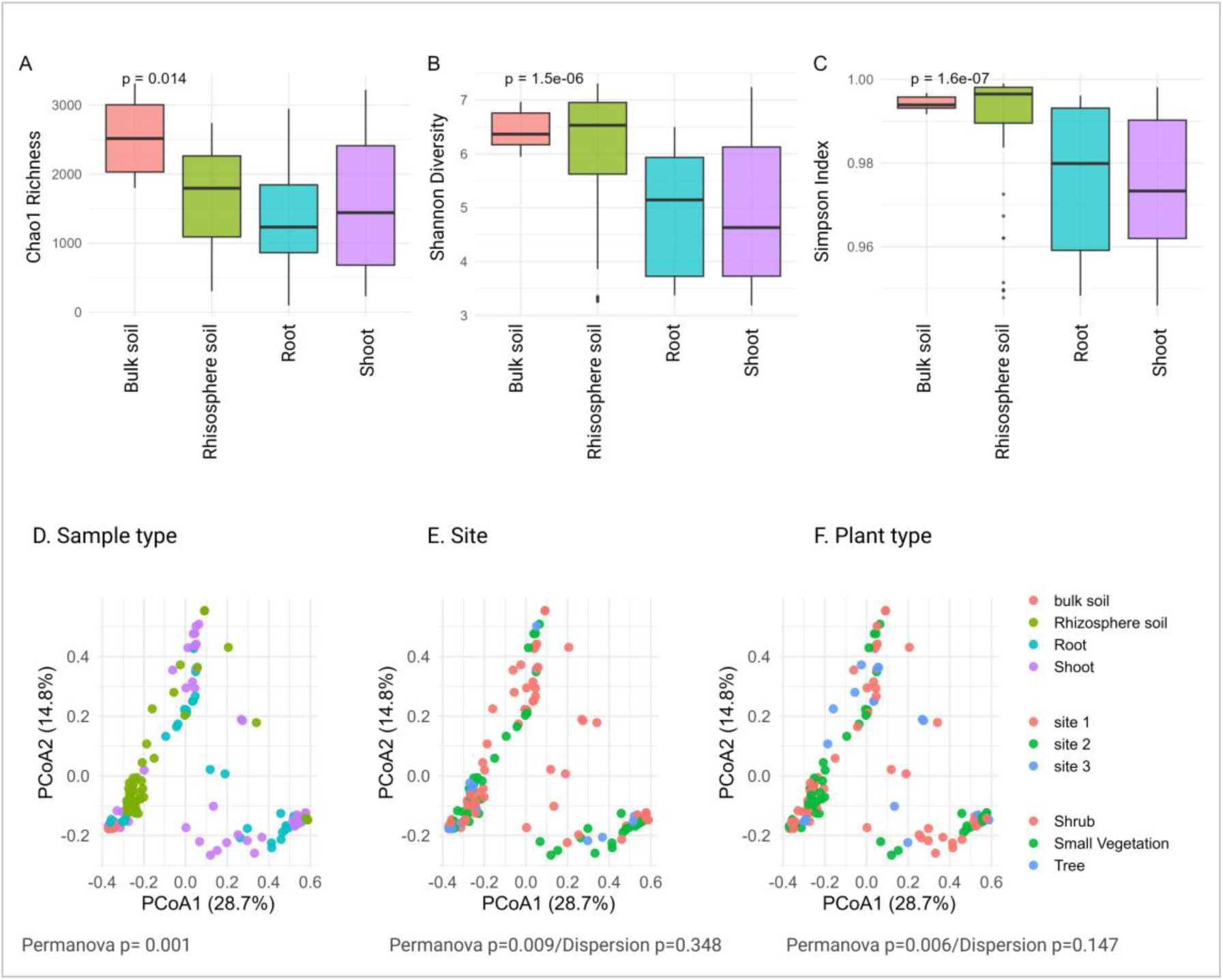
Canonical correspondence analysis (CCA) of bacterial community structure constrained by environmental variables. Arrows represent soil physicochemical parameters, and red labels denote bacterial genera. The direction and length of arrows indicate the strength and direction of each environmental gradient influencing community composition. Environmental factors such as EC, Na₂O, MgO, and CaO showed strong correlations with specific bacterial taxa distributions.

### Culture-dependent isolation and identification of halophytic bacteria

To complement the culture-independent profiling for bacterial communities and validate the presence of dominant bacterial taxa in the sabkha ecosystem, we implemented a culture-dependent approach aimed at isolating of halotolerant bacteria from key halophytes plants, *Juncus acutus* and *Halocnemum strobilaceum*, were selected based on their distinctive signatures of indicator genera (Fig. 4) and core taxa (Fig. S5). Root, shoot, and rhizosphere samples yielded 19 morphologically distinct isolates. Sanger sequencing followed by phylogenetic analysis (Fig. 6) assigned these isolates to several known halophilic or halotolerant taxa, including such *Halomonas*, *Bacillus*, *Salinicola*, *Marinobacter*, *Psychrobacter* and *Marinococcus*. The recovered isolates represent a diverse set of halophilic and halotolerant bacteria associated with sabkha halophytes (Table 1). *Halomonas* isolates (GQ30 and SK63) from *Halocnemum strobilaceum* showed sequence identity (96.4-98.2%) to described halophilic species and tolerated up to 25% NaCl. Additional halophilic taxa included *Idiomarina* sp. MAN K15 and *Marinobacter* sp. PY97S, both isolated from *Halocnemum* shoots. Other isolates, such as *Psychrobacter alimentarius* Psys_23 from *Juncus* roots, a psychrophilic yet halotolerant species, and *Bacillus pumilus* AM2 from *Halocnemum* shoots, are known for broad stress tolerance. Further isolates included *Planomicrobium* sp. XMCCM07 and *Bacillus* sp. TRM82117 from *Juncus* roots, along with an uncultured bacterium and an unidentified strain (DCBB0021) from *Halocnemum*, expanding the phylogenetic breadth of culturable sabkha-associated bacteria.

**Figure 6.**
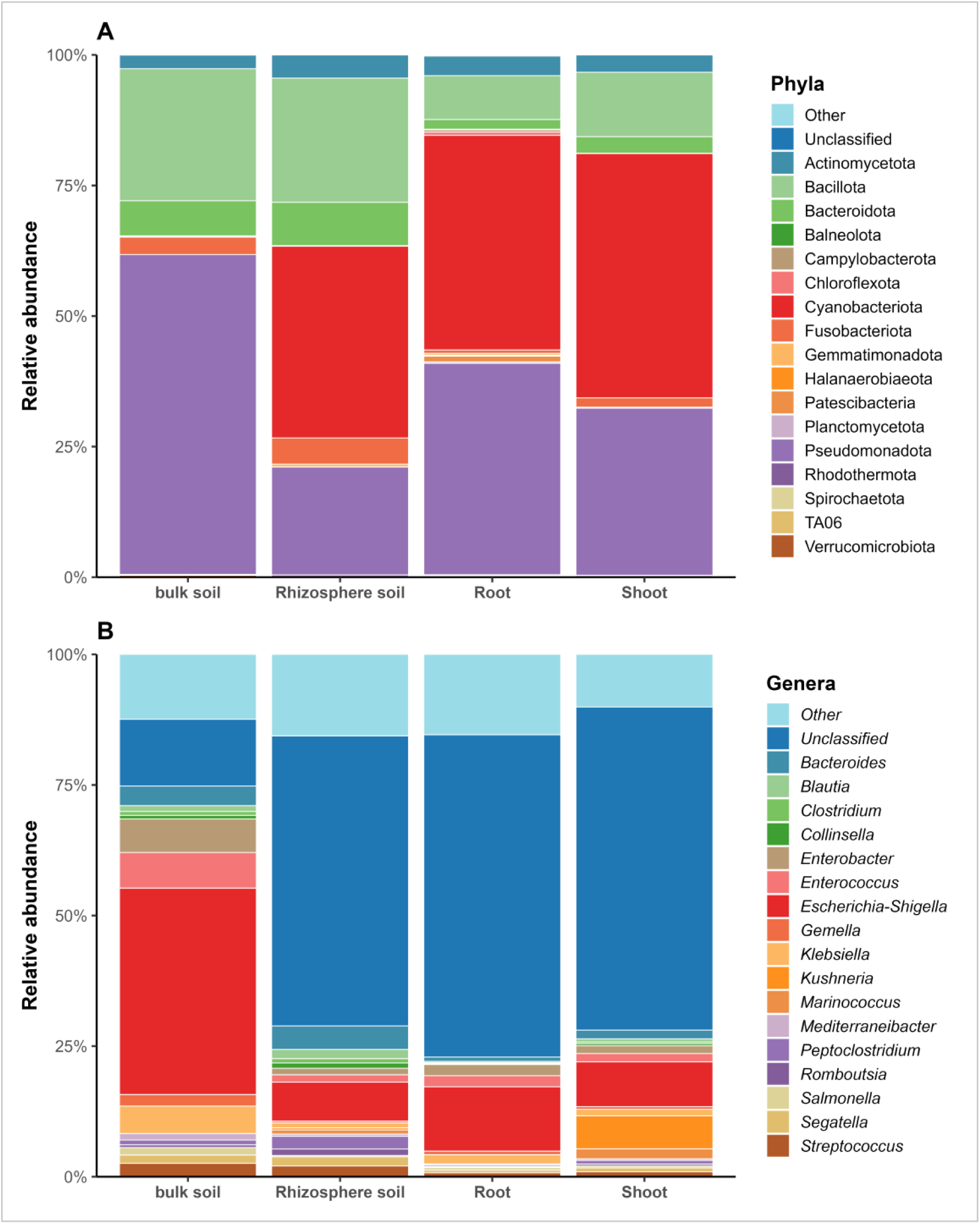
Phylogenetic placement of sabkha bacterial isolates. Maximum-likelihood phylogenetic tree based on 16S rRNA gene sequences showing the relationships between sabkha-derived bacterial isolates (highlighted in red) and reference strains retrieved from the NCBI database. Isolates clustered within well-known halotolerant and halophilic genera, including *Halomonas* spp., *Bacillus* spp., *Salinicola* spp., *Marinobacter* spp., *Psychrobacter* spp., and *Marinococcus* spp. Bootstrap support values greater than 70% are shown at the corresponding nodes.

Phylogenetic analysis revealed that sabkha isolates cluster within major halophilic and stress-adapted lineages (Fig. 6). Many isolates grouped with *Halomonas*, *Idiomarina*, and *Marinobacter*, while others aligned with halotolerant Firmicutes such as *Bacillus* and *Planomicrobium*. Additional isolates branched with *Psychrobacter* and several uncultured taxa, highlighting both known and previously uncharacterized diversity within sabkha microbiomes. Notably, many of these genera overlapped with dominant and indicator taxa identified through sequencing, providing independent confirmation of their prevalence in sabkha environments and demonstrating strong concordance between culture-independent and culture-based approaches. The phylogeny (Fig. 6) underscores the broad taxonomic range and strong enrichment of stress-adapted bacteria associated with halophytes in extreme desert saline ecosystems.

## Discussion

We found that sabkha ecosystems exert strong environmental filtering on bacterial communities, producing highly structured microbiomes across compartments and plant types. Across all sabkhas, pronounced gradients in ionic composition (Na₂O, K₂O), salinity, and pH corresponded to marked shifts in bacterial diversity and composition. We observed a consistent transition from taxonomically rich bulk soil microbiomes toward increasingly selective assemblages within the rhizosphere, root, and shoot, demonstrating that sabkha habitats impose stringent constraints on microbial colonization.

### Environmental filtering dominated bacterial communities

Our findings demonstrate that microbial community assembly in sabkha habitats is overwhelmingly shaped by abiotic forces, supporting our first hypothesis. Extreme salinity, high carbonate content, and a consistently alkaline pH characterised the bulk soils across all three Sabkha sites. These conditions impose substantial selective pressure on microbial physiology [40]. Alpha diversity declined progressively from bulk soil to rhizosphere, root, and shoot compartments, revealing hierarchical filtering along the soil–plant continuum. Beta diversity analyses further supported this trend, showing strong compartmental separation driven by differences in both community centroids and dispersion. Although hierarchical filtering has been widely reported across terrestrial ecosystems [7, 41], the intensity of filtering observed in sabkhas was particularly pronounced, reflecting the extreme environmental constraints shaping microbial community assembly in these saline desert ecosystems.

### Shared halotolerant core microbiota linking hosts and compartments

Despite edaphic heterogeneity among sites, bacterial communities converged on a stable halotolerant core composed primarily of *Halomonas*, *Kushneria*, *Marinococcus* and related genera are well known for osmoprotection, salt tolerance, and metabolic plasticity [42, 43]. This strong cross-site provides clear support for the second hypothesis, indicating deterministic environmental filtering as the major factor shaping microbiome convergence. Remarkably, 66% of ASVs were shared across rhizosphere, root, and shoot compartments, underscoring the persistence of a cross-compartment microbial foundation. The absence of rhizosphere-specific ASVs suggests that the rhizosphere primarily functions as an environmental reservoir from which endophytes are recruited [44]. In contrast, the presence of unique ASVs in root compartments reflects the strongest selective bottleneck along the continuum [45]. Perennial shrubs consistently hosted the richest and most phylogenetically diverse core microbiota across compartments, likely due to their long lifespan, deeper rooting systems, and more stable rhizosphere environments [20, 46].

### Host effects exist but remain secondry to strong environmental filtering

Although plant identity contributed to the structure of sabkha bacterial communities, its influence was modest compared to the dominant effect edaphic filtering, in agreement with the first hypothesis. Across all sites, salinity, ionic composition, and alkalinity exerted the strongest constraints, overshadowing host-driven selection, a pattern widely observed in arid and saline environments [47–50]. Host genotype did produce detectable but subtle signals: for example, *Dichotomicrobium* was enriched in the roots of *Halocnemum strobilaceum*, while indicator taxa such as *Kushneria* and *Halomonas* were associated with *Juncus acutus*, and *Peptoclostridium* with *Frankenia thymifolia*. Similar compartment-specific host effects, more pronounced in the endosphere than in bulk soil or rhizosphere, have been reported in multiple systems, where root exudate chemistry, tissue structure, or plant immune responses shape a subset of microbial taxa [51–54].

However, these host-associated signatures were exceptions rather than the prevailing pattern. The rhizosphere and root microbiota in sabkhas were overwhelmingly structured by abiotic gradients, consistent with global trends in which soil salinity, pH, carbonate content, and aridity explain most variation in microbial community composition, whereas plant identity explains only a minor fraction [55, 56]. Moreover, under extreme environmental stress, such as high ionic strength, osmotic pressure, and desiccation, host-driven selection becomes weaker and more context-dependent, often giving way to stronger deterministic filtering or even stochastic assembly processes [47, 57]. Taken together, our results indicated that while host traits may modulate the abundance of bacterial lineages, salinity and related edaphic factors remain the dominant forces shaping microbiome assembly across sabkha plant species.

### Environmental determinants of bacterial community structure

CCA results further confirmed the central role of edaphic filtering. Salinity-related parameters, particularly EC, Na₂O, K₂O, and carbonate fractions, were the strongest predictors of bacterial community composition. These gradients aligned with halophilic genera such as *Kushneria*, *Halomonas*, and *Marinococcus*, taxa characterized by osmoadaptive mechanisms including compatible solute biosynthesis (ectoine, hydroxyectoine), active transport systems, and ion homeostasis [58, 59]. In contract, copiotrophic taxa such as, *Escherichia–Shigella*, *Enterococcus* occurred in nutrient-enriched microhabitats within high EC. These patterns parallel findings from other hypersaline systems [60], underscoring the global consistency of abiotic filtering under extreme salinity.

### Culture-dependent isolation confirmed stress-adapted bacterial taxa

Culture-dependent analyses provided strong support for the third hypothesis, demonstrating that dominant halophilic taxa detected via metabarcoding are also recoverable through cultivation. Nineteen morphologically distinct isolates were obtained from *J. acutus* and *H. strobilaceum* representing well-known halophilic and halotolerant genera (*Halomonas*, *Idiomarina*, *Marinobacter*, *Psychrobacter*, *Planomicrobium* and *Bacillus*). Several isolates tolerated salinity levels up to 25% NaCl, consistent with established with halophilic physiology [61–63]. Stress-tolerant Bacillota such as *Bacillus pumilus* and *Planomicrobium* were also recovered, consistent with their resilience in desert environments [64]. The correspondence between cultured-independante and culture-based results demonstrates both the ecological dominance and culturable robustness of sabkha halophiles. The isolation of strains clustering with uncultured lineages additionally highlights sabkhas as reservoirs of yet unexplored microbial diversity with promising biotechnological potential.

### Alignment with global patterns of halophyte-assiciated bacteriome

Our findings mirror global patterns observed in halophytic systems. A recent meta-analysis by [65] identified *Thalassospira*, *Erythrobacter,* and *Marinobacter* as predictive halophilic taxa in saline soils, echoing the sabkha core assemblage. Similarly, studies from Utah desert halophytes consistently isolated *Halomonas*, *Kushneria*, and *Bacillus* [66], many of which exhibit extreme salt tolerance (up to 4 M NaCl) and promote plant performance under salinity stress. The strong overlap between our sabkha core microbiome and halophilic lineages reported globally underscores deterministic environmental filtering as a pervasive force shaping plant–microbe associations across saline habitats worldwide.

## Conclusion

This study provides a comprehensive characterization of bacterial communities associated with native halophytes in sabkha ecosystems of southern Morocco. Microbiome assembly in these hypersaline, alkaline habitats is shaped primarily by deterministic environmental filtering, with deep salinity, ionic and carbonate gradients driving a progressive transition from diverse bulk soil microbiota to increasingly selective rhizosphere, root, and shoot communities. Despite site-level heterogeneity, all plant hosts shared a conserved halotolerant core dominated by *Halomonas*, *Kushneria*, and *Marinococcus*, while host identity exerted only secondary influence. The substantial overlap of taxa across compartments highlights the root interior as the most selective microhabitat. Culture-dependent isolation further validated these patterns and recovered culturable representatives of dominant halophilic lineages.

Ecologically, sabkhas emerge as valuable natural laboratories for studying microbial adaptation to extreme abiotic stress. The communities identified here are enriched for traits associated with osmoadaptation, desiccation tolerance, and general stress resilience, providing insight into the mechanisms underpinning survival in hypersaline desert environments. However, the taxonomic and functional resolution of 16S rRNA sequencing limits our ability to identify strain-level adaptive features and specific metabolic pathways. Future metagenomic, metatranscriptomic, and physiological studies will be essential to resolve the genetic basis of halotolerance, especially genes involved in compatible-solute biosynthesis, membrane stabilization, and ion homeostasis. Moreover, although core and indicator taxa were identified, their functional roles remain inferential. Experimental validation through plant inoculation assays and in situ manipulations will be necessary to determine whether these halophilic lineages enhance plant performance under salinity and alkalinity stress. Altogether, the resilient halotolerant taxa highlighted in this study represent promising candidates for the development of targeted microbial inoculants and synthetic consortia (SynComs) aimed at improving crop productivity in saline and alkaline soils. Integrating environmental microbiology with functional genomics and ecological experimentation will pave the way for sustainable biotechnological applications in arid and salt-affected agricultural systems.

## Acknowledgements

We thank Jean Legeay for assistance with high-performance computing data processing and Vittorio Venturi for general support with sequencing. This work was supported by Danilo Licastro at Area Science Park, Trieste, Italy, as part of the Next Generation Sequencing platform. We are grateful to the African Genome Center at Mohammed VI Polytechnic University for providing laboratory facilities. We also thank Mohamed and Tarik at ASARI for their valuable support during field sampling, and ASARI for providing transport and sample storage facilities. Finally, we acknowledge all members of the Phosboucraa project team members for their contributions and support.

## Authors contributions

GA: sampling, experimentation, analysis of sequencing data, and draft the manuscript. NR: designed the sampling strategy, collected the data, data analysis; AEM: sampling, data collection, review manuscript; DL: sequencing and data curation; AH: conceptualization, manuscript review and editing; MH and BA: conceptualization, manuscript drafting, review and supervision. All authors contributed to the article and approved the submitted version.

## Data Availability

The amplicon datasets used in this study are available online with the accession number PRJNA1395314 in the NCBI SRA database.

